# Gene encoding a novel enzyme of LDH2/MDH2 family is lost in plant and animal genomes during transition to land

**DOI:** 10.1101/495713

**Authors:** L.V. Puzakova, M.V. Puzakov, A.A. Soldatov

**Affiliations:** The A.O. Kovalevsky Institute of Marine Biology Research of RAS, Nakhimov av., 2, Sevastopol, Russia, 299011

**Keywords:** evolution, gene loss, oxydoreductase, aquatic enzyme, hypoxic environment, transition to land

## Abstract

L-lactate/malate dehydrogenases (LDH/MDH) and type 2 L-lactate/malate dehydrogenases (LDH2/MDH2) belong to NADH/NADPH-dependent oxidoreductases (anaerobic dehydrogenases). They form a large protein superfamily with multiple enzyme homologs found in all branches of life: from bacteria and archaea to eukaryotes, and play an essential role in metabolism. Here we describe the gene encoding a new enzyme of LDH2/MDH2 oxidoreductase family. This gene is found in genomes of all studied groups/classes of bacteria and fungi. In the plant kingdom this gene was observed only in algae, but not in bryophyta or spermatophyta. This gene is present in all taxonomic groups of animal kingdom beginning with protozoa, but is lost in lungfishes and other, higher taxa of vertebrates (amphibians, reptiles, avians and mammals). Since the gene encoding the new enzyme is found only in taxa associated with the aquatic environment, we named it *AqE (aquatic enzyme)*. We demonstrated that *AqE* gene is convergently lost in different independent lineages of animals and plants. Interestingly, the loss of the gene is consistently associated with transition from aquatic to terrestrial life forms, which suggests that this enzyme is essential in aquatic environment, but redundant or even detrimental in terrestrial organisms.

## Introduction

Water and air environment is characterized by significant differences in physical and chemical conditions. Low solubility of oxygen in water is one of the major factors influencing organisms living in aquatic systems. Oxygen diffusion in water environment is approximately 10000-fold slower than in air (Klyashtorin 1982), causing the development of hypoxic areas in waters with low circulation. Hypoxic zones occur in a wide range of World Ocean aquatic systems (Levin, 2003; Middelburg and Levin, 2009; Gewin, 2010), there concentration of a dissolved oxygen declines below 0.5 mg/l (Levin, 2003; Stoeck et al, 2003). However, large number of aquatic species such as priapulida, loricifera, bivalves, gastropods and benthic fishes adapted for living in hypoxic environment (Levin, 2003; Danovaro et al, 2010), and many hypoxia-tolerant organisms even have evolved to survive at anoxia (Isani et al. 1986; Soldatov et al, 2009; Danovaro et al, 2010). Metabolic pathways in their tissues are substantially reorganized: (a) glycolysis is normally suppressed (Soldatov et al, 2009); (b) transamination of amino acids (glutamate, alanine) is enhanced (Mommsen et al. 1980; Soldatov et al, 2009); (c) mitochondrial electron transport chain reveals non-compensated type of organization (Soldatov and Savina, 2008) etc. Some species (loricifera) do not have typical mitochondria, and their function is performed by relict hydrogenosomas (Danovaro et al, 2010).

“Hypoxic” type of energy metabolism originated earlier during evolution of life and appears to have the widest diversity of metabolic pathways. It is likely that hypoxia-tolerant animals may have undescribed metabolic pathways which function at low-oxygen environment.

Malate-aspartate shuttle is considered to be the most efficient energy producing pathway in hypoxia (anoxia)-tolerant aquatic organisms (Hochachka and Somero, 2002). In anaerobic conditions it allows producing 6 moles of ATP (GTP) per one mole of substrate consumed (Hochachka and Somero, 2002). Cytoplasmic malate dehydrogenase (MDH1, 1.1.1.37) has been suggested to play a key role in malate-aspartate shuttle functioning. MDH1 functioning is associated with aspartate aminotransferase (AST) activity, which catalyzes transamination of asparagine acid (Asp) and α-ketoglutarate into oxaloacetate. Oxaloacetate is reduced by MDH1 to malate with concomitant consumption of glycolytic NADH (Hochachka and Somero 2002):

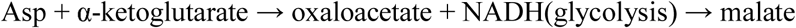

The process described above combines together protein and carbohydrate metabolism. Malate and α-ketoglutarate are transferred into mitochondria through antiport, where they are oxidized to oxaloacetate by mitochondrial MDH2 enzyme. The process is associated with production of NADH. In some mollusks oxaloacetate can be oxidized to propionate allowing respiration chain to produce ATP (GTP) in anaerobic conditions (Hochachka and Somero 2002). Unification of protein and carbohydrate metabolisms is also accomplished by the activity of other malate dehydrogenase family enzymes (for example NADP-malic enzyme, 1.1.1.40) (Hochachka and Somero 2002).

Transition of organisms from water to land has resolved the problem of oxidizer deficiency in aquatic environment, making some metabolic pathways unnecessary. In particular, malate-aspartate shuttle and other anaerobic energy-producing systems have lost their relevance. We hypothesized that in conditions of unlimited oxygen supply “hypoxic” metabolic genes may be lost in genome due to their redundancy for survival. For instance, such selective changes may affect group of oxidoreductase enzymes (MDH and LDH), which enhance the ability of an organism to survive hypoxia.

Here, we describe a gene coding for the new enzyme of LDH2/MDH2 oxidoreductase superfamily. Analysis of the gene ditribution across the tree of life has demonstrated that it is repeatedly (convergently) lost in different independent lineages of animals and plants. Gene sequence was found only in aquatic organisms and was absent in terrestrial animals and plants and, therefore, it was named the *AqE* (*aquatic enzyme*). Since the loss of the gene is consistently associated with transition from aquatic to terrestrial life forms, we speculate that *AqE* enzyme is essential for aquatic environment, but redundant or even detrimental for terrestrial organisms.

## Materials and Methods

Conserved sequences of channel catfish MDH1 (NM_001201146) were analyzed using conserved domain database, CDD (Marchler-Bauer et al. 2017). Prediction structural organization and property of MDH1 amino acid sequences was carried out using the RaptorX software (Källberg et al. 2012; Wang et al. 2016). Multiple alignment of amino acid sequences was performed using MUSCLE (Edgar 2004) with standard settings. Phylogenetic analysis was inferred using the Neighbor-Joining method (Saitou and Nei 1987). The evolutionary distances were computed by the p-distance method (Nei and Kumar 2000). All positions containing gaps and missing data were excluded from the analysis. The analysis of various taxa cytoplasmic malate dehydrogenases (MDH1) involved 28 amino acid sequences of: Acropora digitifera (XP_015758340), Amphimedon queenslandica (XP_019853270), Branchiostoma belcheri (XP_019630870), Hyalella azteca (XP_018014268), Caenorhabditis elegans (CCD71316), Callorhinchus milii (NP_001279824), Ciona intestinalis (XP_018669683), Crassostrea gigas (XP_011449697), Danio rerio (NP_956241), Drosophila melanogaster (NP_609394), Gallus gallus (NP_001303820), Homo sapiens (NP_001303303), Octopus bimaculoides (XP_014784794), Parasteatoda tepidariorum (XP_015918513), Protobothrops mucrosquamatus (XP_015667838), Schistosoma haematobium (XP_012799073), Strongylocentrotus purpuratus (XP_796283), Xenopus tropicalis (NP_001303830), Chelonia mydas (XP_007062950), Chinchilla lanigera (XP_005385779), Bos mutus (XP_005901333), Columba livia (XP_005503807), Falco peregrinus (XP_005232496), Latimeria chalumnae (XP_005989614), Ictalurus punctatus (NP_001188075), Cyprinus carpio (XP_018976734), Trichinella spiralis (XP_003375386), Necator americanus (XP_013301227). The analysis of LDH and MDH enzymes, LDH/MDH-like enzymes and LDH2/MDH2 oxidoreductases involved 25 amino acid sequences of: *Bos taurus* L-lactate dehydrogenase chain A, EC 1.1.1.27 (BC146210); *B. taurus* L-lactate dehydrogenase chain B, EC 1.1.1.27 (BC142006); *B. taurus* L-lactate dehydrogenase chain C, EC 1.1.1.27 (JQ317783); *B. taurus* D-lactate dehydrogenase, EC 1.1.1.28 (BC118204); *Rattus norvegicus* malate dehydrogenase, cytoplasmic, EC 1.1.1.37 (AF093773); *R. norvegicus* malate dehydrogenase, mitochondrial, EC 1.1.1.37 (X04240); *Danio rerio* malate dehydrogenase 1B, EC 1.1.1.-(BC124491); *Mus musculus* NAD-dependent malic enzyme, mitochondrial (malate dehydrogenase (oxaloacetate-decarboxylating), EC 1.1.1.38 (AK042289); *Amaranthus hypochondriacus* NAD-dependent malic enzyme 65 kDa isoform, mitochondrial (malate dehydrogenase (decarboxylating), EC 1.1.1.39 (U01162); *M. musculus* NADP-dependent malic enzyme, mitochondrial (malate dehydrogenase (oxaloacetate-decarboxylating) (NADP+), EC 1.1.1.40 (AK032196); *Arabidopsis thaliana* malate dehydrogenase (NADP+), EC 1.1.1.82 (AY074329); *Escherichia coli* D-malate dehydrogenase (decarboxylating), EC 1.1.1.83 (AAC74870); *E. coli* 3-isopropylmalate dehydrogenase, EC 1.1.1.85 (BAA04537); *E. coli* ureidoglycolate dehydrogenase (4FJU, DOI: 10.2210/pdb4fju/pdb); *Methanocaldococcus jannaschii* (2S)-L-sulfolactate dehydrogenase (2X06, DOI: 10.2210/pdb2x06/pdb); *Phaeobacter gallaeciensis* (2R)-3-sulfolactate dehydrogenase (NADP+), EC 1.1.1.338 (AHD11586); *Pseudomonas syringae* Δ^1^-piperideine-2-carboxylate/Δ^1^-pyrroline-2-carboxylate reductase (2CWF, DOI: 10.2210/pdb2cwf/pdb); *E. coli* 2,3-diketo-L-gulonate reductase (1S20, DOI: 10.2210/pdb1s20/pdb); *Thermus thermophilus* HB8 Type 2 malate/lactate dehydrogenase (1VBI, DOI: 10.2210/pdb1vbi/pdb); *Agrobacterium tumefaciens* malate dehydrogenase (1Z2I, DOI: 10.2210/pdb1z2i/pdb); *Pyrococcus horikoshii* OT3 malate dehydrogenase (1V9N, DOI: 10.2210/pdb1v9n/pdb); *Entamoeba histolytica* malate dehydrogenase (3I0P, DOI: 10.2210/pdb3i0p/pdb); *E. coli* malate/L-lactate dehydrogenases (2G8Y, DOI: 10.2210/pdb2g8y/pdb); *Methanothermus fervidus* malate/(S)-sulfolactate dehydrogenase (WP_013413469); *Ictalurus punctatus* malate dehydrogenase, cytoplasmic (NP_001188075). Evolutionary analysis was performed using MEGA7 software (Kumar et al. 2016). Distribution of AqE gene among taxonomic groups was determined in all NCBI nucleotide and protein databases using BLAST (Altschul et al. 1997; Zhang et al. 2000). Transcriptional activity was assessed by search for homology in TSA and SRA in NCBI databases.

## Results

New LDH2/MDH2 oxidoreductase family enzyme

We have performed a phylogenetic analysis of cytoplasmic malate dehydrogenase (MDH1) amino acid sequences in various taxonomic groups of animals. The channel catfish *Ictalurus punctatus mdh* sequence (NM_001201146) forms a distinct branch of phylogenetic tree (Figure 1). Multiple alignment of amino acid sequences demonstrated substantial difference of *I. punctatus mdh* gene comparing to the sequences from other species. In particular, clear distinctions were noticed in the most conservative fragments of the *I. punctatus mdh* gene, which were similar in other studied species. This raises the question whether channel catfish *mdh* gene indeed encodes cytoplasmic malate dehydrogenase.

**Figure 1.**
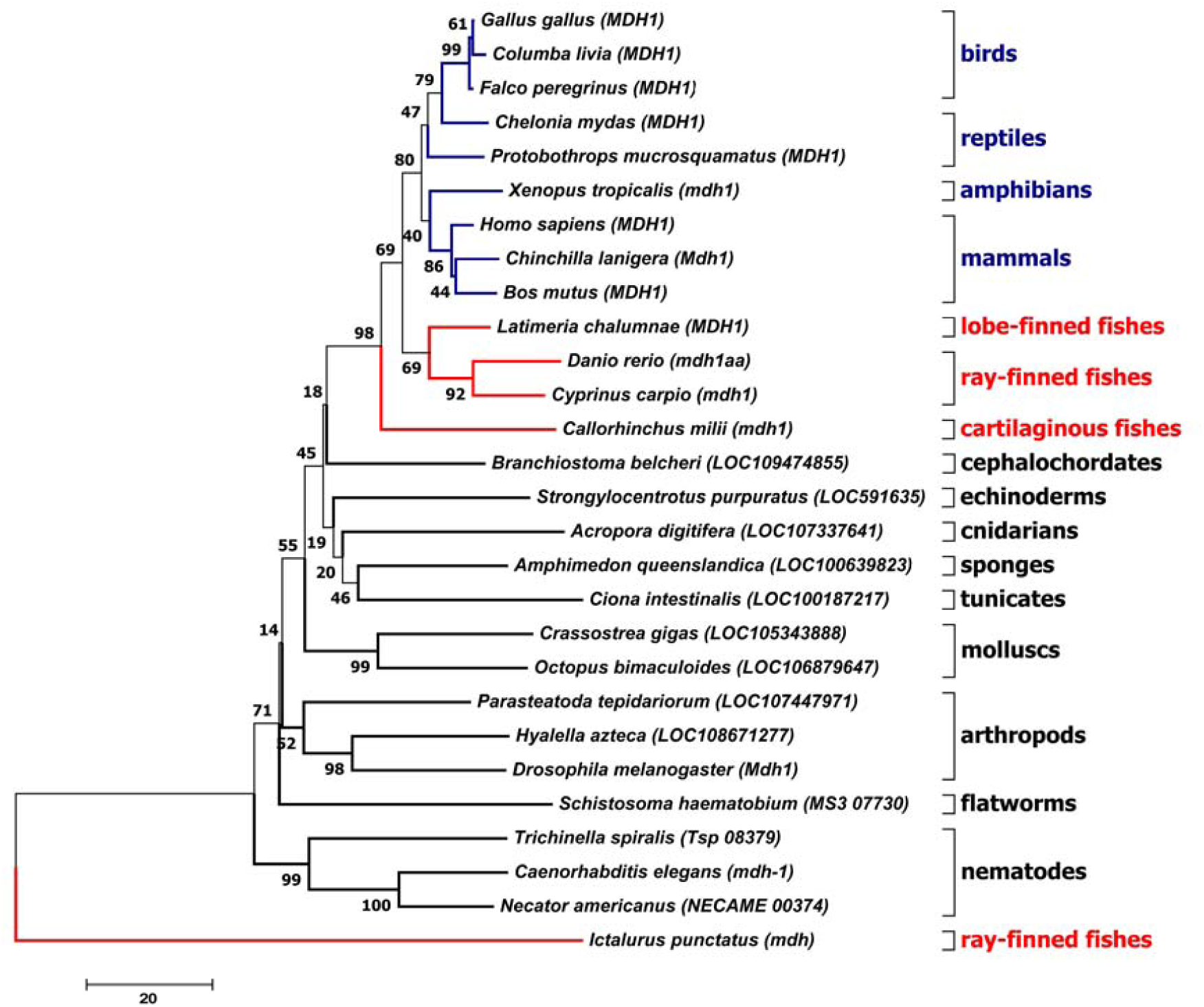
Evolutionary relationships of cytoplasmic malate dehydrogenases (MDH1) in various taxa.

Examination of the *I. punctatus mdh* sequence in conservative domain database (CDD) (Marchler-Bauer et al. 2017) demonstrated that the protein contains conservative domain Ldh_2 (pfam02615) which is usually present in enzymes of in LDH/MDH superfamily. This superfamily contains NAD(P)-dependent dehydrogenase, which utilize lactate and malate as a substrate (Honka et al. 1990).

The evolutionary history was inferred using the Neighbor-Joining method, bootstrap 1000. The analysis involved cytoplasmic malate dehydrogenases amino acid sequences of 28 organisms (in parentheses is the gene name). Evolutionary analyses were conducted in MEGA7 (Kumar et al, 2016). The tetrapodas taxa is indicated by blue color. The fishes taxa is indicated by red color.

Prediction of structure and properties of the discovered enzyme identified the highest homology to 3I0P protein (P-value: 2.11e-11, uGDT(GDT): 268(74), Score: 382), 2X06:A protein (P-value: 6.26e-11, uGDT(GDT): 268(74), Score: 366) and 1V9N protein (P-value: 1.56e-10, uGDT(GDT): 263(73), Score: 352). Although tertiary structure of 3I0P protein (malate dehydrogenase from *Entamoeba histolytica*, DOI: 10.2210/pdb3i0p/pdb) and 1V9N protein (malate dehydrogenase from *Pyrococcus horikoshii* OT3, DOI: 10.2210/pdb1v9n/pdb) has been already determined, their cellular functions remain unknown. They were classified as members of MDH family solely on the basis of sequence homology. The 2X06 protein (sulfolactate dehydrogenase from *Methanocaldococcus jannaschii*, DOI: 10.2210/pdb2x06/pdb) is a member of LDH2/MDH2 oxidoreductase family and utilizes L-2-hydroxycarboxylates such as (2R)-3-sulfolactate, (S)-malate, (S)-lactate, and (S)-2-hydroxyglutarate) (Graupner et al. 2000; Irimia et al. 2004). Thus, there is no strong reason to state that the *I. punctatus mdh* is cytoplasmic malate dehydrogenase gene.

We have used phylogenetic analysis to define relation of the channel catfish protein (which was annotated as MDH) to know enzyme families. The analysis included canonical LDH and MDH enzymes, LDH/MDH-like enzymes and LDH2/MDH2 oxidoreductases with known function as well as those enzymes which properties were predicted by sequence homology. The *I. punctatus* MDH amino acid sequence from *I. punctatus* groups with L-sulfolactate dehydrogenase-like enzymes clade (Figure 2), indicating that this protein is not cytoplasmic malate dehydrogenase, but rather novel enzyme.

**Figure 2.**
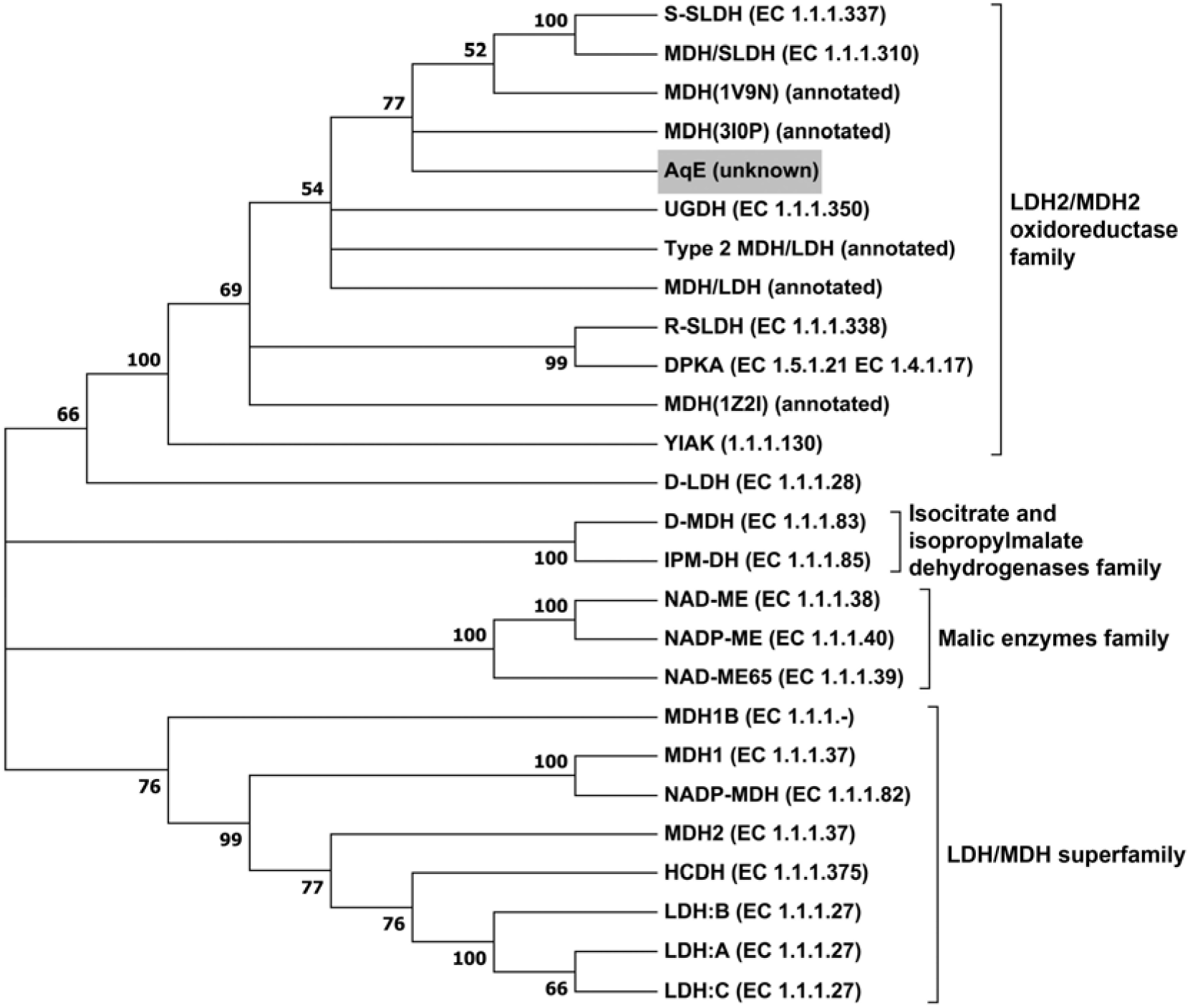
Phylogenetic analysis of LDH and MDH enzymes, LDH/MDH-like enzymes and LDH2/MDH2 oxidoreductases.

Multiple alignment of amino acid sequences were created by MUSCLE (Edgar 2004) and a Neighbor-Joining tree was built using MEGA7 (Kumar et al, 2016). The analysis involved 25 amino acid sequences: LDH:A - *Bos taurus* L-lactate dehydrogenase chain A; LDH:B - *B. taurus* L-lactate dehydrogenase chain B; LDH:C - *B. taurus* L-lactate dehydrogenase chain C; D-LDH -*B. taurus* D-lactate dehydrogenase; MDH1 - *Rattus norvegicus* malate dehydrogenase, cytoplasmic; MDH2 - *R. norvegicus* malate dehydrogenase, mitochondrial; MDH1B - *Danio rerio* malate dehydrogenase 1B; NAD-ME – *Mus musculus* NAD-dependent malic enzyme, mitochondrial (malate dehydrogenase (oxaloacetate-decarboxylating); NAD-ME65 - *Amaranthus hypochondriacus* NAD-dependent malic enzyme 65 kDa isoform, mitochondrial (malate dehydrogenase (decarboxylating); NADP-ME - *M. musculus* NADP-dependent malic enzyme, mitochondrial (malate dehydrogenase (oxaloacetate-decarboxylating) (NADP+); NADP-MDH - *Arabidopsis thaliana* malate dehydrogenase (NADP+); D-MDH - *Escherichia coli* D-malate dehydrogenase (decarboxylating); IPM-DH - *E. coli* 3-isopropylmalate dehydrogenase; UGDH - *E. coli* ureidoglycolate dehydrogenase; S-SLDH - *Methanocaldococcus jannaschii* (2S)-L-sulfolactate dehydrogenase; R-SLDH - *Phaeobacter gallaeciensis* (2R)-3- sulfolactate dehydrogenase (NADP+); DPKA - *Pseudomonas syringae* Δ1-piperideine-2- carboxylate/Δ1-pyrroline-2-carboxylate reductase; YIAK - *E. coli* 2,3-diketo-L-gulonate reductase; Type 2 MDH/LDH - *Thermus thermophilus* HB8 Type 2 malate/lactate dehydrogenase; MDH(1Z2I) - *Agrobacterium tumefaciens* malate dehydrogenase; MDH(1V9N) - *Pyrococcus horikoshii* OT3 malate dehydrogenase; MDH(3I0P) - *Entamoeba histolytica* malate dehydrogenase; MDH/LDH - *E. coli* malate/L-lactate dehydrogenases; MDH/SLDH - *Methanothermus fervidus* malate/(S)-sulfolactate dehydrogenase; AqE - *Ictalurus punctatus* malatedehydrogenase.

## Distribution of *AqE* among living organisms

It was interesting to determine distribution of the new enzyme among different life forms. Gene sequence was found only in aquatic organisms and was absent in terrestrial animals and plants and, therefore, it was named the *AqE* (*aquatic enzyme*). Searching in NCBI nucleotide and protein sequence databases has shown that *AqE* existed in fungi, protozoa and bacteria (Figure 3). In the plants homologous sequence was found only in algae, which are primitive lower plants inhabiting mostly aquatic environments. In higher (or terrestrial) plants the gene coding for this enzyme was not observed in species from briophyta to angiosperms. In animals the protein existed in ctenophora, sponges, cnidarians, mollusks, worms, arthropods, echinodermata, amphioxus and tunicates. *AqE* gene was also found in teleosts and chondrichthyes. In lobe-finned fishes *AqE* gene existed in *Latimeria chalumnae*, a member of coelacanths. However, gene sequence of this species is divided in two parts in the database. Two exons of the gene are located in scaffold 04923 (NW_005823933) and the rest seven are present in scaffold 02115 (NW_005821125). Moreover, methionine start codon is absent in nucleotide sequence. This observation suggests that *L. chalumnae AqE* gene sequence may have changed by the insertion of transposable elements or dramatic rearrangement.

**Figure 3.**
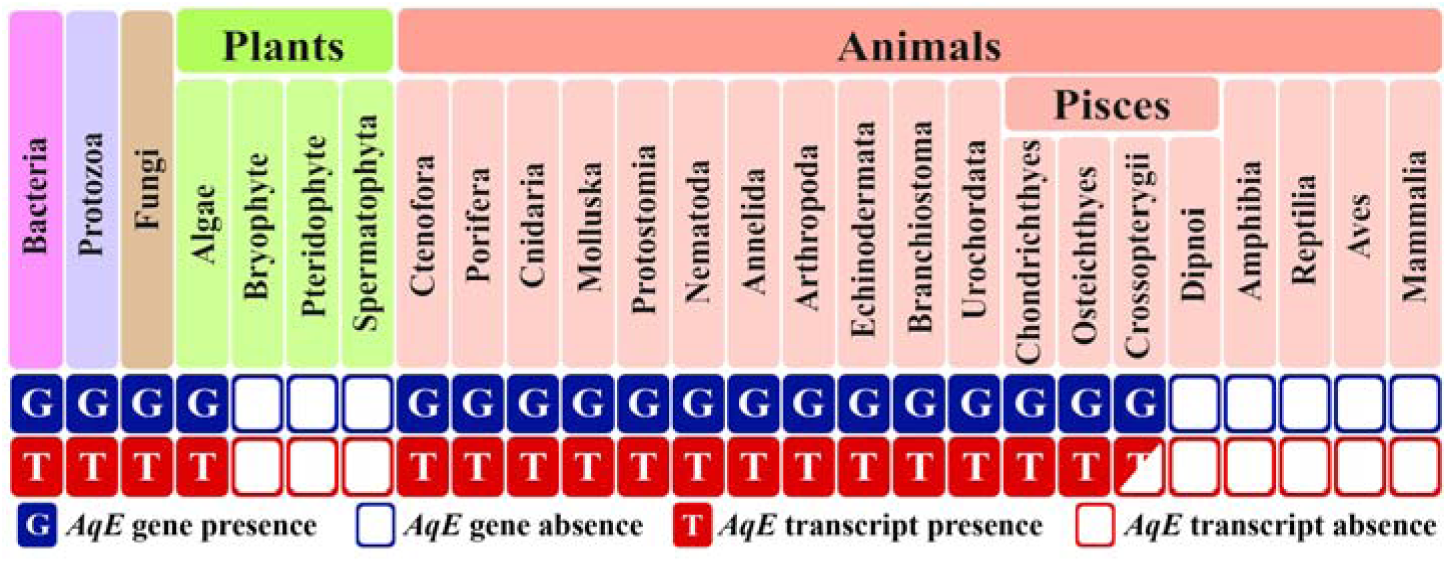
Taxonomic distribution and expression of *AqE* gene across the distinct organism groups.

In taxonomic group of actinistians, lungfishes, which are the closest living relative of tetrapods, *AqE* gene was not observed.

An analysis of genomic and transcriptomic databases of the tetrapods revealed several dozens of fragments homologous to *AqE* sequence. Each of these sequences from the tetrapods was analyzed by *blastn* and *blastx* (Supplementary table 1). They were shown to have a higher degree of homology with the *AqE* sequences from the bacteria, fungi, nematodes, and insects, than with *AqE* sequence of the channel catfish (NM_001201146). Due to evolutionary relationship the homology between the sequences from the fish and the tetrapods is expected to be higher, than that between the tetrapods and the more distantly related life forms (insects, nematodes or prokaryotes). For example, the fragments found in the TSA database of the Chinese salamander, *Hynobius chinensis*, demonstrate a high degree of homology (85-87%) to the insect *AqE* gene, but do not have homologous sequences in the genome this amphibian, probably, reflecting a contamination of amphibian DNA samples by the insect DNA. Insects are the main food resource for amphibians, providing plausible source for contamination of samples used for genome sequencing.

*AqE* sequence found in the TSA database of a reptile, *Anolis carolinensis*, shows 100% homology to the sequences from the cichlides, while no homology was found among the WGS sequences of *A. carolinensis*. The homologous *AqE* fragments found in mammals show the higher degree of sequence similarity to the fungal, bacterial, or archaeal sequences, than to *AqE* of the channel catfish (see Online resource 1).

Thus, all *AqE* homologous sequences from the tetrapods, most likely represent a contamination by the genomes of lower organisms. Moreover, *AqE* homologs were detected simultaneously in genomes and in transcriptomes of the organisms belonging to the groups, in which the aquatic enzyme was found. In contrast, in tetrapods homologies were, as a rule, found only in either genomic or in transcriptomic sequences, but not simultaneously in both. Thus, we concluded that amphibians, reptiles, birds and mammals, as well as the terrestrial plants do not possess the sequences encoding aquatic enzyme.

## Analysis of AqE gene activity

To determine wherever the *AqE* gene is transcribed into corresponding RNA we performed analysis of its transcription activity in all taxonomic groups containing *AqE* full-length sequence. Every group of organisms was examined using expression data obtained from TSA. We established that *AqE* is expressed in all organisms containing this gene except of lobe-finned fishes (latimeria) (Figure 3). The *L. chalumnae* transcriptome has not been fully assembled yet, and it contains reads exhibiting only weak homology to the *AqE* sequence. We assume that the absence of homologous for AqE reads in the latimeria could result from the loss of gene functionality. Since, *AqE* gene is transcribed in all organisms containing its sequence, is it likely that it takes part in metabolic processes.

## Discussion

Our analysis identified a convergent loss of *AqE* gene in two utterly distinct evolutionary branches (plants and animals), during their transition from water to land. This fact suggests a non-accidental nature of the process. Which factor might cause the loss of gene involved in metabolism? It is possible that alteration of living conditions (and corresponding selective pressures) appears to be the most obvious reason causing metabolism reorganization making *AqE* gene dispensable for organism survival. One of the key difference between aquatic and air environment is the oxygen concentration. Even in the oxygen-saturated water it’s concentration does not exceed 10 ml/l, which is 21 times lower than in the air. Therefore, breathing conditions is much more complicated for aquatic animals, than for air-breathing organisms. Aquatic organisms survive in conditions of periodic or long-termed hypoxia (anoxia) by acquiring various adaptive strategies including anaerobic metabolism. In contrast, atmosphere is characterized by constant abundant level of oxygen. Thus, it is reasonable to hypothesize that atmospheric oxygen concentration, might play a key role in loss of *AqE*.

Several scenarios have been proposed to explain the process of evolutionary gene inactivation or loss: (a) slow accumulation of mutations in gene resulting in transformation to pseudogene and gradual degradation (fragmentation); (b) accidental gene loss (deletion) due to unequal crossing over in meiosis or mobile genetic elements transposition (Albalat and Cañestro 2016). The deletion appears to be more parsimonious explanation of the AqE gene loss, since no homology to *AqE* gene and its fragments was detected in the genomes of higher plants and terrestrial animals. In plant kingdom *AqE* gene loss occurred on the frontier between algae and bryophytes. Algae are primitive lower plants, which predominantly inhabit aquatic environments. In contrast, bryophytes, which utilize atmospheric oxygen, belong to terrestrial higher plants with relatively complicated anatomy. In animal kingdom gene loss most likely occurred in the relict lobe-finned fishes, which are the closest tetrapod ancestors. Our analysis showed that latimeria most likely possesses non-functional *AqE* gene or only its part is expressed. In lung-fishes and all taxonomic groups of tetrapods the gene is fully excluded (Figure 3).

Gene loss is a widely spread phenomena in evolution. The intensive genomic data acquisition has unexpectedly demonstrated that the gene loss may not only be evolutionally neutral but also can substantially increase adaptive potential of the specie. For example, Greenberg et al (2003) suggested that loss of desaturase 2 (*Desat2*) activity in *D. melanogaster* lines is accompanied by increased tolerance to colder environment. Similarly, excluding of genes encoding several group of chemoreceptors in fruit fly resulted in changes in food preference and adaptations to new ecological niches (Clark et al. 2007; McBride et al. 2007; Goldman-Huertas et al. 2015).

Likewise, it was suggested, that human myosin gene MYH16 has lost functionality after modification of a food diet (Stedman et al. 2004). This fact, in turn, caused reduction of the importance of jaw muscles during early hominoid evolution. It was proposed that MYH16 gene loss could have led to skull volume increase and growth of the brain during human evolution. Mutations leading to loss of gene functionality seems to occur more frequently than those leading to acquisition of new features (Albalat and Cañestro 2016). It is, therefore, plausible that gene loss mutations have a greater impact on adaptive evolution, especially during rapid adjustment to the abrupt environmental changes.

Alternatively, a gene may become redundant, and in this case its loss is neutral effect on vital function. For instance, various vertebrates (teleost fishes, passeriform birds, bats, guinea pigs and anthropoid primates) independently have lost L-gulonolactone oxidase (GLO) gene which encodes protein involved in late stages of vitamin C biosynthesis. This loss was made possible as a result of replacing the low-vitamin C diet to rich-vitamin food ration (Moreau and Dabrowski 1998; Drouin et al. 2011).

*AqE* gene has been lost independently in two distinct taxonomic groups and this loss coincided with transition from water to land. Such a powerful factor as atmospheric oxygen, rather than others, could have an evolutionary effect on the cell metabolic processes. A unique feature of LDH2/MDH2 oxidoreductases is its ability to utilize large number of substrates for oxidation, including Δ^1^-piperideine-2-carboxylate/Δ^1^-pyrroline-2-carboxylate, 2,3-diketo-L-gulonate, L-sulfolactate, malate, lactate, ureidoglycolate. *L*-sulfolactate is the most common substrate for L-sulfolactate dehydrogenase, that provides cells with additional source of reducing agents used for respiration (Denger and Cook 2010):

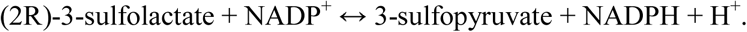

L-sulfolactate is produced in the sediments as a result of archaeas (*Methanocaldococcus sp)* methanogenic activity (Irimia et al. 2004). It can be speculated that water contains certain concentration of dissolved L-sulfolactate, which may be used as an applicable substrate for oxidation by aquatic organisms. However, this metabolic pathway is utterly redundant for terrestrial organisms. In this case loss of the gene exhibiting homology with sulpholactate dehydrogenase enzymes becomes understandable. However, terrestrial animals which spend a portion of their life-cycle in aquatic saprotrophic environment, such as insects, should form an exception to this rule. Thus, our analyses identified that insects retained the transcriptionally active *AqE* gene, which is in agreement with our main hypothesis. The same logic can be apply to explain why fungi also preserved *AqE* gene.

If we assume that deletion of *AqE* is evolutionary neutral, then it remains unclear how this deletion became wide-spread in population of ancestral tetrapods. Also, this hypothesis fails to explain why species with defective *AqE* gene has not been found in our research. It is possible that *AqE* gene loss in both plants and animals could have occurred randomly and was independently pickup in evolution due to founder effect. However, an adaptive scenario seems more plausible to explain a convergent pattern of *AqE* gene loss. Thus, if deletion of *AqE* gene provides an advantage in atmospheric environment the resulting selective pressure would force a convergent fixation of null allele in populations of animals and plants. Definitive conclusions about adaptive significance of *AqE* gene loss could be drawn after determining enzyme function(s).

## Supporting information

## Acknowledgments

Under support of the Russian Academy of Sciences research grant № AAAA-A18-118021490093-4

