## Supplementary material for "Gene encoding a novel enzyme of LDH2/MDH2 family is lost in plant and animal genomes during transition to land"

**Table 1 Search results for tetrapoda's AqE in the NCBI databases**

| homologous AqE sequence | | | | | check result in blastn (nucleotide) | | | | | | check result in blastx (proteins) | | | | | |
|---|---|---|---|---|---|---|---|---|---|---|---|---|---|---|---|---|
| **Species** | **Accession number** | **Query cover** | **E value** | **Ident** | **Species** | **Nucleotide sequence** | **Query cover** | **E value** | **Ident** | **Accession number** | **Species** | **Protein** | **Query cover** | **E value** | **Ident** | **Accession number** |
| Amphibia (WGS) | | | | | | | | | | | | | | | | |
| No significant similarity found | | | | | | | | | | | | | | | | |
| Amphibia (TSA) | | | | | | | | | | | | | | | | |
| Hynobius chinensis | GAQK01015698 | 67% | 1,00E-071 | 47% | Stomoxys calcitrans | PREDICTED: uncharacterized oxidoreductase YjmC (LOC106092848), transcript variant X4, mRNA | 71% | 4,00E-071 | 71% | XM_013259779 | Chironomus tentans | Larval EST Library Chironomus tentans cDNA, mRNA sequence | 86% | **0,0** | 87% | JZ928071 |
| Hynobius chinensis | GAQK01057269 | 54% | 3,00E-056 | 48% | Drosophila busckii | PREDICTED: uncharacterized oxidoreductase YjmC-like (LOC108600592), mRNA | 63% | 8,00E-047 | 71% | XM_017988270 | Clunio marinus | CLUMA_CG015751, isoform A | 98% | **2,00E-140** | 85% | CRL02920 |
| Hynobius chinensis | GAQK01059480 | 20% | 8,00E-017 | 50% | Phlebotomus papatasi | BAC clone PLPAAEX-22H12 from chromosome unknown | 61% | 1,00E-012 | 73% | AC238782 | Clunio marinus | CLUMA_CG015751, isoform A | 98% | **2,00E-140** | 85% | CRL02920 |
| Hynobius chinensis | GAQK01052735 | 21% | 1,00E-010 | 35% | Pediculus humanus corporis | malate dehydrogenase, putative, mRNA | 84% | 5,00E-012 | 71% | XM_002428328 | Clunio marinus | CLUMA_CG015751, isoform A | 98% | **2,00E-140** | 85% | CRL02920 |
| Oreobates cruralis | GFNJ01017475 | 70% | 5,00E-078 | 47% | Brugia malayi | Malate/L-lactate dehydrogenase family protein partial mRNA | 59% | 1,00E-085 | 69% | XM_001893344 | Toxocara canis | L-sulfolactate dehydrogenase | 64% | **2,00E-166** | 78% | KHN83282 |
| Oreobates cruralis | GFNJ01055057 | 51% | 1,00E-060 | 52% | Necator americanus | malate/L-lactate dehydrogenase mRNA | 93% | 2,00E-054 | 69% | XM_013448284 | Toxocara canis | Malate dehydrogenase | 94% | **2,00E-115** | 81% | KHN76802 |
| Oreobates cruralis | GFNJ01157730 | 30% | 6,00E-028 | 48% | Caenorhabditis elegans | Uncharacterized protein (F36A2.3), partial mRNA | 52% | 1,00E-028 | 77% | NM_059977 | Caenorhabditis brenneri | hypothetical protein CAEBREN_26413 | 97% | **6,00E-047** | 78% | EGT39622 |
| Oreobates cruralis | GFNJ01114607 | 18% | 3,00E-016 | 51% | Ascaris lumbricoides | WGS scaffold0000270 | 44% | 6,00E-014 | 76% | LK872126 | Toxocara canis | L-sulfolactate dehydrogenase | 60% | **6,00E-039** | 73% | KHN83282 |
| Oreobates cruralis | GFNJ01115729 | 14% | 2,00E-010 | 57% | Ascaris lumbricoides | genome assembly A_lumbricoides_Ecuador_v1_5_4, scaffold ALUE_scaffold0000270 | 44% | 3,00E-012 | 76% | LK872126 | Toxocara canis | L-sulfolactate dehydrogenase | 60% | **6,00E-025** | 68% | KHN83282 |
| Oreobates cruralis | GFNJ01171472 | 19% | 2,00E-009 | 46% | Haemonchus placei | genome assembly H_placei_MHpl1, scaffold HPLM_scaffold0001238 | 23% | 6,00E-013 | 84% | LM584294 | Toxocara canis | Malate dehydrogenase | 57% | **6,00E-020** | 73% | KHN76802 |
| Rana catesbeiana | LH280374 | 14% | 1,5 | 33% | Entamoeba nuttalli | malate dehydrogenase, putative partial mRNA | 73% | 1,00E-026 | 75% | XM_008861456 | Kosmotoga | MULTISPECIES: lactate dehydrogenase | 96% | **3,00E-033** | 66% | WP_012744996 |
| Reptilia (WGS) | | | | | | | | | | | | | | | | |
| No significant similarity found | | | | | | | | | | | | | | | | |
| Reptilia (TSA) | | | | | | | | | | | | | | | | |
| Anolis carolinensis | GBDE01259384 | 17% | 8,00E-015 | 49% | Haplochromis burtoni | PREDICTED: uncharacterized oxidoreductase YjmC-like (LOC102301235), mRNA | 100% | **6,00E-139** | 100% | XM_014335344 | Haplochromis burtoni | PREDICTED: LOW QUALITY PROTEIN: uncharacterized oxidoreductase YjmC-like | 69% | 1,00E-040 | 100% | XP_014190830 |
| Anolis carolinensis | GAFZ01259162 | 17% | 8,00E-015 | 49% | Haplochromis burtoni | PREDICTED: uncharacterized oxidoreductase YjmC-like (LOC102301235), mRNA | 100% | **6,00E-139** | 100% | XM_014335344 | Haplochromis burtoni | PREDICTED: LOW QUALITY PROTEIN: uncharacterized oxidoreductase YjmC-like | 69% | 1,00E-040 | 100% | XP_014190830 |
| Thamnophis sirtalis | GDKU01003202 | 83% | 5,00E-103 | 47% | Caenorhabditis briggsae | Hypothetical protein CBG03139 (CBG03139) mRNA | 90% | 9,00E-086 | 69% | XM_002631271 | Caenorhabditis latens | hypothetical protein FL83_09348 | 99% | **1,00E-160** | 74% | OZG16426 |
| Thamnophis sirtalis | GDKU01003203 | 63% | 1,00E-077 | 47% | Caenorhabditis elegans | Uncharacterized protein (VF13D12L.3), partial mRNA | 85% | 6,00E-075 | 70% | NM_064099 | Caenorhabditis briggsae | Hypothetical protein CBG03139 | 95% | **2,00E-127** | 74% | XP_002631317 |
| Thamnophis sirtalis | GDKU01125775 | 58% | 2,00E-059 | 45% | Necator americanus | malate/L-lactate dehydrogenase mRNA | 86% | 3,00E-072 | 70% | XM_013448284 | Ancylostoma ceylanicum | malate/L-lactate dehydrogenase | 95% | **9,00E-142** | 79% | EPB69155 |
| Thamnophis sirtalis | GDKU01125776 | 54% | 6,00E-053 | 45% | Necator americanus | malate/L-lactate dehydrogenase mRNA | 91% | 2,00E-072 | 70% | XM_013448284 | Ancylostoma ceylanicum | malate/L-lactate dehydrogenase | 94% | **4,00E-134** | 81% | EPB69155 |
| Thamnophis sirtalis | GDKU01125774 | 33% | 1,00E-036 | 53% | Necator americanus | malate/L-lactate dehydrogenase mRNA | 76% | 2,00E-058 | 75% | XM_013448284 | Necator americanus | malate/L-lactate dehydrogenase | 91% | **2,00E-077** | 75% | XP_013303738 |
| Aves (WGS) | | | | | | | | | | | | | | | | |
| No significant similarity found | | | | | | | | | | | | | | | | |
| Aves (TSA) | | | | | | | | | | | | | | | | |
| No significant similarity found | | | | | | | | | | | | | | | | |
| Mammalia (WGS) | | | | | | | | | | | | | | | | |
| Alces americanus | FLNZ01006080 | 85% | 2,00E-040 | 32% | Methanobrevibacter olleyae | complete genome | 99% | **0,0** | 78% | CP014265 | Methanobrevibacter ruminantium M1 | L-sulfolactate dehydrogenase ComC | 44% | **0,0** | 87% | ADC47830 |
| Alces americanus | FLNY01002128 | 95% | 5,00E-035 | 30% | Clostridium saccharolyticum | complete genome | 48% | 3,00E-061 | 71% | CP002109 | Clostridiales bacterium | lactate dehydrogenase | 100% | **0,0** | 84% | OKZ47964 |
| Alces americanus | FLNW01003395 | 91% | 2,00E-032 | 31% | Clostridium saccharolyticum | complete genome | 53% | 1,00E-054 | 70% | CP002109 | Clostridiales bacterium | lactate dehydrogenase | 100% | **0,0** | 81% | OKZ47964 |
| Alces americanus | FLNW01002402 | 88% | 1,00E-031 | 29% | No significant similarity found. | | | | | | Betaproteobacteria bacterium | hypothetical protein A3G25_09970 | 99% | **5,00E-098** | 43% | OGA54328 |
| Alces americanus | FLNZ01001500 | 88% | 1,00E-031 | 29% | No significant similarity found. | | | | | | Betaproteobacteria bacterium | hypothetical protein A3G25_09970 | 41% | **5,00E-102** | 43% | OGA54328 |

| | | | | | | | | | | | | | | | | |
|---|---|---|---|---|---|---|---|---|---|---|---|---|---|---|---|---|
| Ammotragus lervia | NIVO01071597 | 79% | 1,00E-020 | 26% | Pseudomonas sp. | complete genome | 64% | 4,00E-139 | 75% | CP015992 | Microvirga vignae | oxidoreductase | 52% | 1,00E-128 | 60% | WP_047188095 |
| Capreolus capreolus | CCMK013088418 | 92% | 5,00E-029 | 29% | Pseudomonas libanensis | complete genome | 100% | 0,0 | 91% | LT629699 | Pseudomonas sp. | lactate dehydrogenase | 94% | 0,0 | 99% | WP_065952525 |
| Capreolus capreolus | CCMK010257545 | 92% | 7,00E-028 | 28% | Pseudomonas antarctica | complete genome | 99% | 0,0 | 84% | LT629704 | Pseudomonas | MULTISPECIES: lactate dehydrogenase | 100% | 0,0 | 97% | WP_065931262 |
| Capreolus capreolus | CCMK013088109 | 79% | 9,00E-027 | 29% | Pseudomonas libanensis | complete genome | 98% | 0,0 | 85% | LT629699 | Pseudomonas | MULTISPECIES: lactate dehydrogenase | 100% | 0,0 | 100% | WP_065931262 |
| Capreolus capreolus | CCMK012998051 | 52% | 4,00E-018 | 33% | Pseudomonas fluorescens | complete genome | 99% | 0,0 | 88% | CP015638 | Pseudomonas fluorescens | lactate dehydrogenase | 87% | 1,00E-130 | 98% | WP_069075835 |
| Homo sapiens | OCRZ01000383 | 85% | 3,00E-033 | 31% | Clostridium stercorarium subsp. Leptospartum | complete genome | 12% | 6,00E-099 | 69% | CP014673 | Sphingobacteriia bacterium | type I restriction endonuclease subunit M | 20% | 0,0 | 87% | OHC88889 |
| Homo sapiens | CQU01000275 | 92% | 6,00E-029 | 27% | Carnobacterium sp. | complete genome | 37% | 9,00E-037 | 69% | CP002563 | Longilinea arvoryzae | lactate dehydrogenase | 99% | 7,00E-154 | 57% | WP_075073005 |
| Homo sapiens | LWKW01189682 | 21% | 1,00E-014 | 47% | Paenibacillus sabinae | complete genome | 100% | 4,00E-026 | 72% | CP004078 | Paenisporosarcina sp. | hypothetical protein | 96% | 1,00E-030 | 66% | WP_017382155 |
| Homo sapiens | CYGL01005383 | 60% | 1,00E-013 | 27% | Escherichia coli | complete genome | 100% | 0,0 | 100% | CP022466 | Escherichia coli | 3-dehydro-L-gulonate 2-dehydrogenase | 42% | 1,00E-178 | 100% | WP_097755041 |
| Lipotes vexillifer | AUPI01006348 | 57% | 2,00E-010 | 29% | Pseudomonas sp. | complete genome | 100% | 0,0 | 99% | LN865164 | Pseudomonas putida | Delta(1)-pyrroline-2-carboxylate/Delta(1)-piperideine-2-carboxylate reductase | 100% | 3,00E-129 | 100% | WP_023535133 |
| Mus musculus | AAHY01767366 | 96% | 4,00E-040 | 30% | Neurospora crassa | actin-like protein arp-6, variant (NCU05587), mRNA | 100% | 0,0 | 100% | XM_011396669 | Neurospora crassa | malate/L-lactate dehydrogenase, variant | 99% | 0,0 | 100% | XP_011394969 |
| Mus musculus | AAHY01773846 | 38% | 3,00E-015 | 33% | Escherichia coli strain | complete genome | 100% | 0,0 | 100% | CP024859 | uncultured bacterium | malate/L-lactate dehydrogenase | 99% | 9,00E-098 | 100% | AMJ35043 |
| Mus musculus | AAHY01773368 | 15% | 1,9 | 35% | Escherichia coli | complete genome | 100% | 0,0 | 99% | CP024859 | Escherichia coli | ureidoglycolate dehydrogenase | 36% | 7,00E-065 | 100% | WP_074460053 |
| Pantholops hodgsonii | AGTT01240857 | 94% | 3,00E-026 | 29% | Agrobacterium rhizogenes | complete genome | 99% | 0,0 | 98% | CP019702 | Agrobacterium rhizogenes | oxidoreductase | 100% | 0,0 | 99% | WP_065117039 |
| Pantholops hodgsonii | AGTT01264776 | 79% | 2,00E-020 | 26% | Rhizobium sp | complete genome | 58% | 4,00E-084 | 75% | CP021371 | Microvirga vignae | oxidoreductase | 98% | 9,00E-125 | 62% | WP_047188095 |
| Pantholops hodgsonii | AGTT01261918 | 97% | 3,00E-018 | 26% | Sphingobium yanoikuyae | complete genome | 100% | 0,0 | 99% | CP023741 | Sphingobium yanoikuyae | oxidoreductase | 93% | 0,0 | 100% | WP_097382989 |
| Pantholops hodgsonii | AGTT01244910 | 68% | 9,00E-018 | 24% | Agrobacterium rhizogenes | complete genome | 100% | 0,0 | 99% | CP019701 | Agrobacterium rhizogenes | sulfolactate dehydrogenase | 100% | 7,00E-173 | 100% | WP_065114557 |
| Pantholops hodgsonii | AGTT01244926 | 93% | 1,00E-013 | 26% | Agrobacterium rhizogenes | complete genome | 100% | 0,0 | 99% | CP019702 | Agrobacterium rhizogenes | lactate dehydrogenase | 100% | 0,0 | 99% | WP_065117926 |
| Pteronotus parnellii | PPAWGZ01437054 | 70% | 4,00E-031 | 33% | Fusarium oxysporum f. sp. Lycopersici | alcohol dehydrogenase mRNA | 100% | 0,0 | 99% | XM_018379891 | Fusarium oxysporum f. sp. Vasinfectum | alcohol dehydrogenase | 100% | 0,0 | 100% | EXM24453 |
| Sorex araneus | AALT02200735 | 8% | 1,00E-025 | 32% | Parastrongyloides trichosuri | complete genome | 19% | 7,00E-015 | 78% | LM523163 | Trichuris suis | malate/L-lactate dehydrogenase | 89% | 3,00E-053 | 53% | KHJ48592 |
| Sus scrofa | AJKK01174601 | 56% | 1,00E-022 | 34% | Reinekea forsetii | complete genome | 13% | 7,00E-015 | 70% | CP011797 | Pseudomonas | MULTISPECIES: hypothetical protein | 57% | 2,00E-170 | 95% | WP_073451712 |
| Sus scrofa | AJKK01160500 | 92% | 3,00E-015 | 25% | Pseudomonas sp. | complete genome | 100% | 0,0 | 99% | LN865164 | Pseudomonas | MULTISPECIES: lactate dehydrogenase | 100% | 0,0 | 100% | WP_027914861 |
| Sus scrofa | AJKK01074756 | 25% | 3,00E-011 | 42% | Pseudomonas sp. | complete genome | 100% | 2,00E-119 | 97% | LT629744 | Pseudomonas | MULTISPECIES: lactate dehydrogenase | 99% | 2,00E-051 | 100% | WP_054897479 |
| Mammalia (non-reduntant) | | | | | | | | | | | | | | | | |
| Sorex araneus | XM_012936024 | 63% | 6,00E-073 | 48% | Trichinella spiralis | malate dehydrogenase (Tsp_11783) mRNA, complete cds | 71% | 6,00E-042 | 69% | XM_003374148 | Trichuris suis | malate/L-lactate dehydrogenase | 99% | 5,00E-110 | 71% | KHJ48592 |
| Mammalia (EST) | | | | | | | | | | | | | | | | |
| Mus musculus | Z31186 | 19% | 5,00E-006 | 37% | Escherichia coli | complete genome | 100% | 6,00E-087 | 100% | CP024859 | Shigella sonnei | malate/lactate/ureidoglycolate dehydrogenase | 78% | 1,00E-039 | 100% | WP_052979015 |
| Sus scrofa | EV959923 | 57% | 9,00E-019 | 28% | Escherichia coli | complete genome | 97% | 0,0 | 99% | CP024859 | Escherichia coli | malate/lactate/ureidoglycolate dehydrogenase | 96% | 1,00E-153 | 99% | WP_089722270 |
| Sus scrofa | AJ747487 | 33% | 1,00E-004 | 27% | Escherichia coli | complete genome | 98% | 0,0 | 99% | CP024859 | Escherichia coli | ureidoglycolate dehydrogenase | 50% | 6,00E-083 | 100% | WP_074460053 |
| Mammalia (TSA) | | | | | | | | | | | | | | | | |
| Tupaia chinensis | JU169505 | 69% | 2,00E-022 | 28% | Escherichia coli | complete genome | 100% | 0,0 | 100% | CP024859 | Escherichia coli | malate/lactate/ureidoglycolate dehydrogenase | 99% | 0,0 | 100% | WP_089602881 |
